# Study-Phase Reinstatement

**DOI:** 10.1101/2023.10.04.560946

**Authors:** David J. Halpern, Bradley C. Lega, Robert E. Gross, Chengyuan Wu, Michael R. Sperling, Joshua P. Aronson, Barbara C. Jobst, Michael J. Kahana

**Affiliations:** Dept. of Psychology, Univ. of Pennsylvania, 425 S. Univ. Ave., Philadelphia, 19104, PA, USA; Dept. of Neurosurgery, Univ. of Texas Southwestern, Dallas, TX, USA; Dept. of Neurosurgery, Emory School of Medicine, Atlanta, GA, USA; Dept. of Neurosurgery, Thomas Jefferson Univ., Philadelphia, PA, USA; Dept. of Neurology, Thomas Jefferson Univ., Philadelphia, PA, USA; Dept. of Neurosurgery, Beth Israel Deaconess Med. Ctr., Boston, MA, USA; Dept. of Neurology, Dartmouth-Hitchcock Med. Ctr., Lebanon, NH, USA

**Keywords:** Memory, Reinstatement, intracranial EEG, Psychology, Neuroscience

## Abstract

Can the brain improve the retrievability of an experience after it has occurred? Systems consolidation theory proposes that item-specific cortical reactivation during post-encoding rest periods facilitates the formation of stable memory representations, a prediction supported by neural evidence in humans and animals. Such reactivation may also occur on shorter time scales, offering a potential account of classic list memory phenomena but lacking in support from neural data. Leveraging the high-temporal specificity of intracranial electroencephalography (iEEG), we investigate spontaneous reactivation of previously experienced items during brief intervals between individual encoding events. Across two large-scale free recall experiments, we show that reactivation during these periods, measured by spectral iEEG similarity, predicts subsequent recall. In a third experiment, we show that the same methodology can identify post-encoding reactivation that correlates with subsequent memory, consistent with previous results. Thus, spontaneous study-phase reinstatement reliably predicts memory behavior, linking psychological accounts to neural mechanisms and providing evidence for rapid consolidation processes during encoding.

## 1 Introduction

Recent evidence from functional magnetic resonance imaging (fMRI) and intracranial electroencephalography (iEEG) suggests that spontaneous neural reactivation of specific previously encoded content during sleep or awake rest predicts subsequent item recognition [1], cued recall [2, 3] and reconstruction [4] of that content. This is explained theoretically by neuroscientific two-stage models of memory [5–7], which propose that reactivation during post-encoding rest periods [8, 9] strengthens memory by consolidating fragile hippocampal memories into stable cortical representations. Consolidation occur spontaneously, even, or especially, without cues from the external environment.

Similarly, psychological theories of study-phase retrieval [e.g. 10–14] and data on overt rehearsal [e.g. 15–17] also propose that mental reactivation during or between encoding other experiences also determines subsequent recall probability and organization. The psychological construct of rehearsal is likely implemented by *cortical* reactivation of the relevant neural representations. In a recent study, Bird and colleagues [18] instructed participants to rehearse previously seen videos when cued while in an fMRI scanner. The degree of reinstatement during these rehearsal periods correlated with subsequent memory for video details. This work, along with similar results showing that (lack of) scalp EEG reinstatement underlies directed forgetting [19], suggests a very similar mechanism for cued rehearsal and *spontaneous* consolidation. The overlap invites speculation that consolidation-like processes may happen more frequently and on shorter time scales than previously studied.

While some theorists [20, 21] consider the possibility of more frequent or opportunistic consolidation periods, the consensus remains that sleep and rest are special times for consolidation [6, 7, 22] and thus the existing neural evidence for reactivation focuses on these time periods. Recent work suggests that the brain can transition rapidly between “online” externally-driven states and “offline” consolidation-promoting states on the time-scale of seconds [23, 24] but no study to date has linked these states to content reinstatement. It, therefore, remains unknown whether rapid, *spontaneous*, item-specific, study-phase neural reinstatement predicts subsequent recall, much as reinstatement during post-encoding periods does. Furthermore, it is unclear whether neural measurements of reinstatement relate to in memory in a similar way as the psychological construct of rehearsal. Studies of overt rehearsal [15] show that rehearsal not only correlates with recall probability but also predicts output order (more rehearsal predicts earlier recall) and organization (items rehearsed together are also recalled together). The present work leverages the high temporal precision of intracranial EEG and three large-scale free recall experiments to address these knowledge gaps.

Further studies demonstrating that neural pattern similarity of repeated items predicts subsequent memory for those items [25, 26]. The author’s interpretation of this result is that such measures index covert study-phase retrieval of previous presentations. Psychologists have invoked study phase retrieval to explain the spacing effect [27–29], wherein increased spacing of repeated items leads to better recall [12, 13, 30]. Feng and colleagues [31] demonstrated that increased neural similarity occurs when comparing an item to its own repeated presentation, as opposed to a different item, but this effect is seen only with spaced repetitions rather than massed repetitions. This suggests that neural similarity plays a role in the observed behavioral spacing effect. However, reinstatement in these studies involving repeated items resulted from the stimulus presentation rather than occurring spontaneously.

In perhaps the most direct evidence for spontaneous study-phase reinstatement, Wu and colleagues [32, 33] investigated how reactivation during a delay, after viewing a sequence of images, affected subsequent memory for sequence details. In one experiment with semantically coherent sequences, they found that the degree of reinstatement of image-related neural activity differed between sequences where subjects recalled an above-or below-average number of details in a subsequent cued recall test. In contrast, there was no effect for non-coherent sequences, leading Wu and colleagues to conclude that people reinstate sequences during study to extract higher-level semantic information. Unlike the current study, Wu and colleagues did not assess whether the reinstatement of image-related activity improved memory for that *specific* image,. Thus, the reinstatement they found could represent a reinstatement of the entire list context, leaving open the question of whether item-specific study-phase reinstatement predicts future memory.

Our work asks whether spontaneous reinstatement during a study phase predicts future recall, as suggested by psychological explanations of memory behavior. These theories suggest that, in addition to externally-driven perceptual experiences, internally-driven thoughts, reminders, retrieved memories and imagined scenarios can themselves be encoded in memory, greatly expanding the role of neural reactivation beyond prior studies. We use two large, independent data sets with intracranial EEG to relate the degree of spontaneous item-specific reinstatement while encoding other experiences (study-phase reinstatement) to subsequent recall probability and organization for that item. In a separate third dataset using the same methodological approach, we ask whether we observe similar reinstatement during an unfilled delay interval between the initial encoding period and later recall, as in previous reactivation studies. Overall, these analyses aim to determine if rapid, spontaneous, study-phase reinstatement plays a role in memory akin to post-encoding consolidation.

## 2 Results

All three data sets involved intracranial EEG recorded during variations on free recall tasks (see Figures 1 and 4). Subjects studied twelve words and, following a brief delay, attempted to recall as many as they could remember. In Experiments 1 (N = 215) and 2 (N = 257), subjects performed a math distractor task during the delay interval; in Experiment 3 (N = 48), the interval remained unfilled. In Experiment 3, some words repeated two or three times in the same list, resulting in 27 word presentations in total during the encoding interval. In Experiment 1, words on the list were drawn from three semantic categories while in the other two experiments, words were drawn at random from a wordpool. Due to a programming error, encoding presentations were slightly shorter in Experiment 3 than in Experiments 1 and 2. In all other respects, the three experiments did not differ, providing the opportunity to replicate our key results several times. Neurosurgical patients performed all three experiments while intracranial EEG was being recorded. These neural recordings provide the basis for identifying latent spontaneous reinstatement

**Fig. 1.**
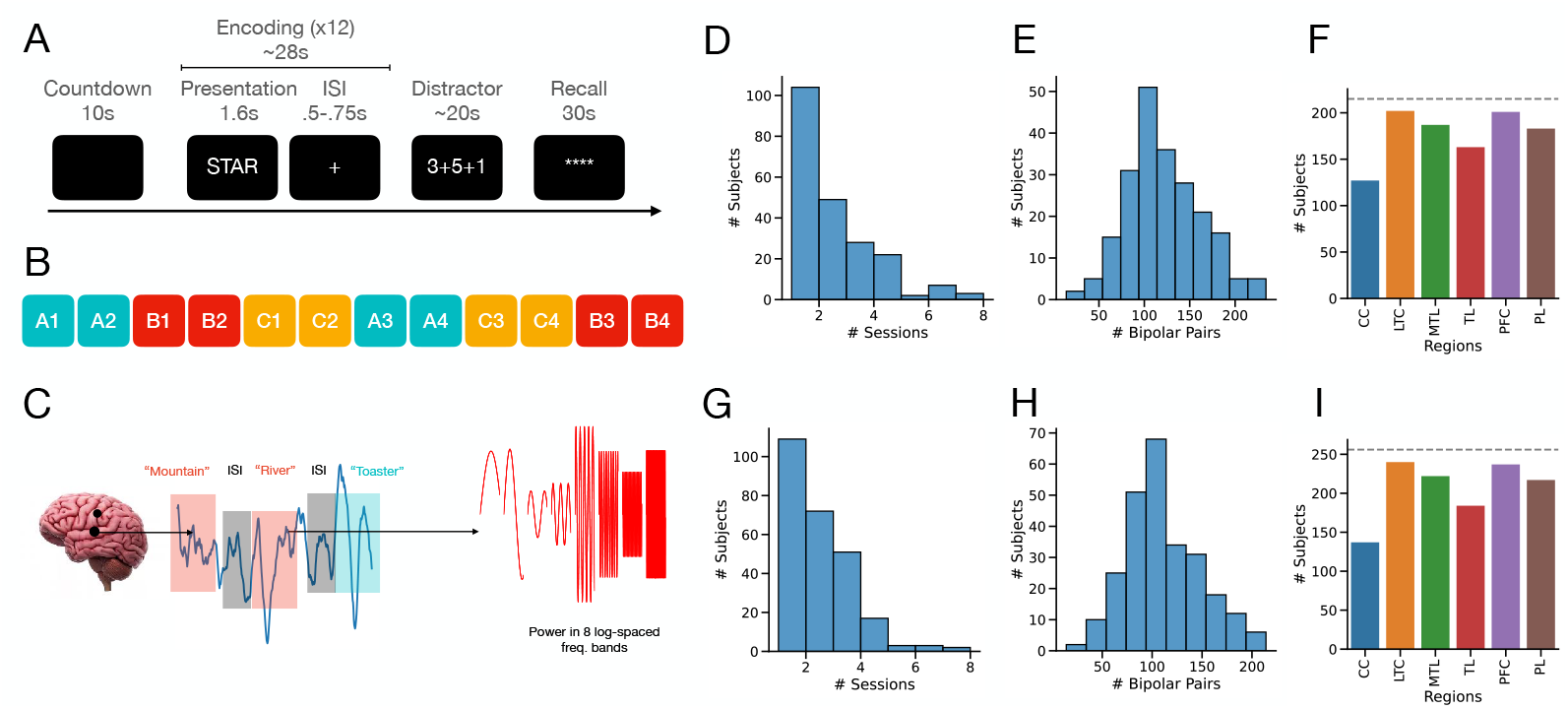
Experimental tasks and methods for Experiments 1 and 2. A. Timeline for a single trial of free recall. Following a ten second countdown, twelve words appear sequentially on screen for 1.6 seconds each with .5 to .75 second inter-stimulus intervals. Following this encoding period, subjects perform twenty second math distractor task in order to prevent rehearsal. Finally, during the recall period, subjects have 30 seconds to recall as many words as possible. B. Category structure of Experiment 1 on a given twelve word list. Four words are drawn from three categories and presented in a block structure as shown. C. Methods used for constructing spectral features. EEG signals are taken each electrode from specific time period, e.g., while a word is on the screen or during an inter-stimulus interval, and then decomposed into power in 8 log-spaced frequency bands. They are then compared to this spectral representation at other time points, e.g., during the inter-stimulus interval. D. Number of sessions collected per subject in Experiment 1. E. Number of bipolar pairs per subject in Experiment F. Number of subjects with bipolar pairs in each region for Experiment 1. (*CC* = Cingulate Cortex, *LTC* = Lateral Temporal Cortex, *MTL* = Medial Temporal Lobe, *TL* = (Other) Temporal Lobe, *PFC* = Prefrontal Cortex, *PL* = Parietal Lobe) G. Number of sessions collected per subject in Experiment 2. H. Number of bipolar pairs per subject in Experiment 2. I. Number of subjects with bipolar pairs in each region for Experiment 2.

We used a spectral representational similarity analysis (RSA) [34–37] approach to studying reinstatement. This method analyzes the predictors of the similarity between frequency representations of neural signals at two time points. We first decomposed the neural signals from each bipolar pair of electrodes into power in eight log-spaced frequency bands. We then compute the z-transformed [38] Pearson correlation between the signals at the two time points across all electrodes and frequencies. This correlation measure is our dependent metric of reinstatement in the following analyses.

### 2.1 Semantic pattern similarity

Our main analyses, aimed at detecting item reactivation during inter-stimulus intervals, rely on the neural patterns representing the semantic content of the items. We therefore begin by validating our RSA-based [34, 35] approach by examining the degree to which time and semantic content determined spectral power similarity during the encoding periods in Experiment 1, which involved categorized lists. By comparing the similarity of items from the same category to items from different categories, we can determine whether semantic content drives spectral pattern similarity. Figure 2 shows the correlation between the spectral power representations at two encoding time points on the same list as a function of their absolute serial position difference and whether or not the words come from the same semantic category. The dominant effect on pattern similarity is distance in time (as represented by an item’s serial position). However, items from the same category are more similar than items from different categories across all serial position distances, suggesting that semantics additionally determines a component of neural pattern similarity. These analyses conceptually replicate effects demonstrated in work by Manning and colleagues [36, 37], using a different data set and method. In addition, Kragel and colleagues [39] demonstrated the ability to predict serial position and semantic category using the same spectral power components in a subset of 69 subjects from the current dataset. However, neither of those papers demonstrates these two effects simultaneously in the same neural signals, as shown here. Finally, a key difference between our analyses and past work using intracranial EEG (including the papers previously mentioned as well as [1]) is that we did not perform any selection of electrodes or time points prior to analyzing pattern similarity. To test the semantic effect, we fit a mixed effect model predicting the pattern similarity between signals associated with two items as a function of whether the items were from the same category and the temporal distance between the items in terms of serial position. We found significant main effects of serial position distance (*F* (1, 92.26) = 153.89, *p* < .001) and the two items being from the same category ((*F* (1, 5.83) = 19.15, *p* < .01). We did not find a significant interaction between the two (*F* (1, 22.14) = 2.87, *p* = .1). Examining the point estimates, we found that, as expected, similarity decreases as a function of temporal distance and increases if the items are from the same category.

**Fig. 2.**
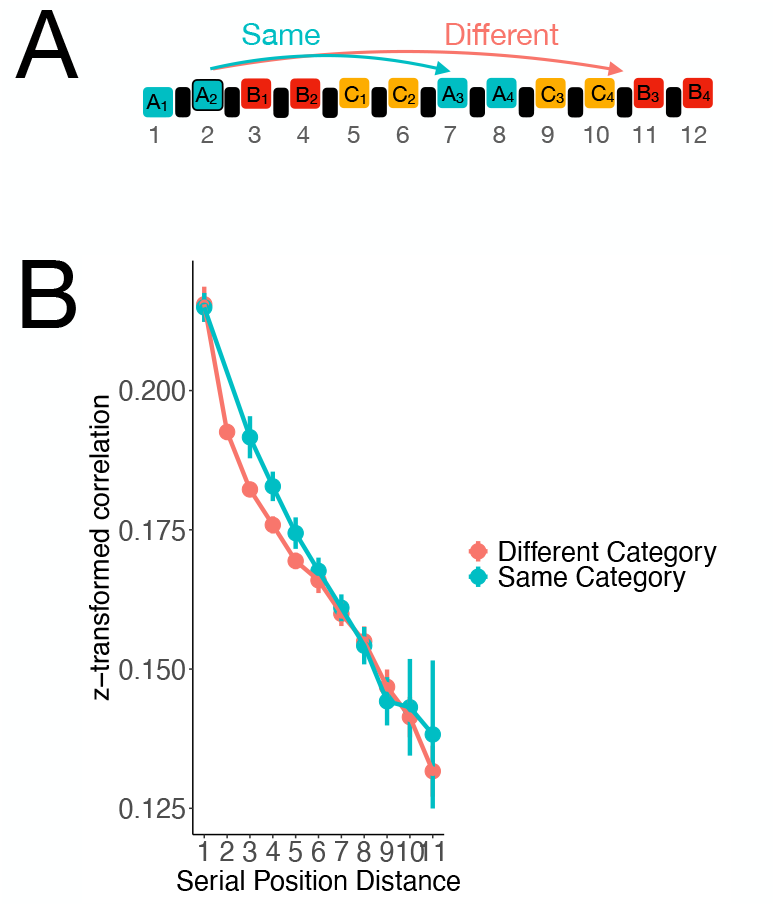
Here, we investigate the similarity of spectral representations of neural signals during two word presentations on the same list. We plot similarity as a function of the absolute serial position distance between the two items and whether they came from the same category or not. Because of the constraints on list order in the experimental design, words from the same category could not appear at a distance of two serial positions away. A. Schematic demonstrating the list structure and which items are involved in each set of comparisons in Figure B. B. Similarity computed during encoding presentations Experiment 1 (N = 215). Error bars are 95% confidence intervals and reflect variation within list using an approach based on Cousineau [40].

### 2.2 Study-phase reinstatement

Having validated the pattern similarity measure during encoding and replicated prior results, we now use the same measure to examine study-phase reinstatement. For this analysis, we investigate whether neural patterns associated with a particular encoded item reactivate during the inter-stimulus intervals following future list items. We used a mixed effects model to compare patterns of spectral power between item presentations (during the 750-1000ms ISI period, the *ISI*) with power while a word was presented on the screen (the *initial encoding presentation*). We predict similarity as a function of whether the initially presented word was itself subsequently recalled, whether the word prior to the ISI was recalled and whether the two words were from the same category. We include these variables in the analysis because we would expect that both of these variables would increase similarity between the initial encoding presentation and the ISI even without spontaneous reinstatement. We additionally examined whether reinstatement differed as a function of recall organization in two ways, based on results from the overt rehearsal literature [15]. First, we would expect a greater degree of reinstatement for the first recalled item than items recalled later. Second, we would expect that the degree of reinstatement during a particular ISI would predict the strength of associations between the item before the ISI and the reinstated item and therefore the likelihood that items will later be recalled sequentially. We used the following specification for the fixed effects (with the full model described in the *Methods* section):

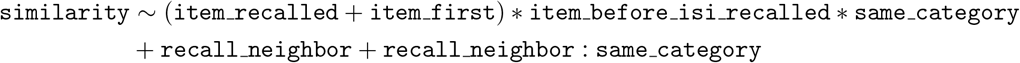

`item recalled` and `item first` reflect whether the specific item was recalled and whether it was recalled first. `item before isi recalled` indicates whether the item before the ISI was recalled and `same category` indicates that the initial encoding item and the item before the ISI are drawn from the same category. Finally, `recall neighbor` indicates whether the encoding item and the item before the ISI were recalled next to each other during the recall period, given that both items were recalled. For Experiment 2, we used the same model but without the terms involving `same category` because items in this Experiment were randomly sampled from a common wordpool. Random effects in the model ensure that all relevant comparisons are *within-list*.

Given the results from our first analysis, it is likely that the effects of recency and categorical similarity drive much of pattern similarity. To focus on spontaneous reinstatement, we therefore limit the analyses to pattern similarity comparisons involving initial encoding presentations from the first half of the list (the first six serial positions) and ISIs from the latter half (the five intervals between the last six serial positions). In the case of Experiment 1, we also ensured that there was at least one category pair between the initial encoded item and the comparison ISI. This ensures that any observed reactivation is unlikely to be due to recency and similarity alone.

In Experiment 1, we observed a main effect of `item recalled` such that items that were subsequently recalled were more similar to activity during subsequent ISIs during encoding than items that were not subsequently recalled (Fig. 3B, (*F* (1, 13.4) = 10.39, *p* = .006)). We do not find reliable evidence that this effect varies by the recall status of the item before the ISI (*F* (1, 28.6) = 2.47, *p* = .13) or whether the item before the ISI was from the same category (*F* (1, 16.0) = .66, *p* = .43). Consistent with this finding, post-hoc comparisons revealed significant effects of recall status, regardless of whether the item before the ISI was from the same category or was itself recalled (marginally significant in the case of different category, item-before-ISI not recalled, *ps* = [.007 − .09]). We could not reliably identify whether the items recalled first had consistently different levels of reactivation than other recalled items (*F* (1, 67.4) = 67.4, *p* = .84). We also did not see consistent differences between items that were sequentially recalled and pairs of items that were both recalled but not sequentially (*F* (1, 17.0) = .07, *p* = .8). Additionally, we find that the recall status of the item before the ISI is a consistently positive predictor of similarity (*F* (1, 11.2) = 6.33, *p* = .03), indicating that some component of the similarity may be due to cognitive processes that occur when viewing items that subjects subsequently recall. However, two items being from the same category is not a significant predictor (*F* (1, 24.6) = .663, *p* = .4), although, as can be seen in Fig. 3B, the estimates are slightly higher when the two items are from the same category, the direction we would predict.

**Fig. 3.**
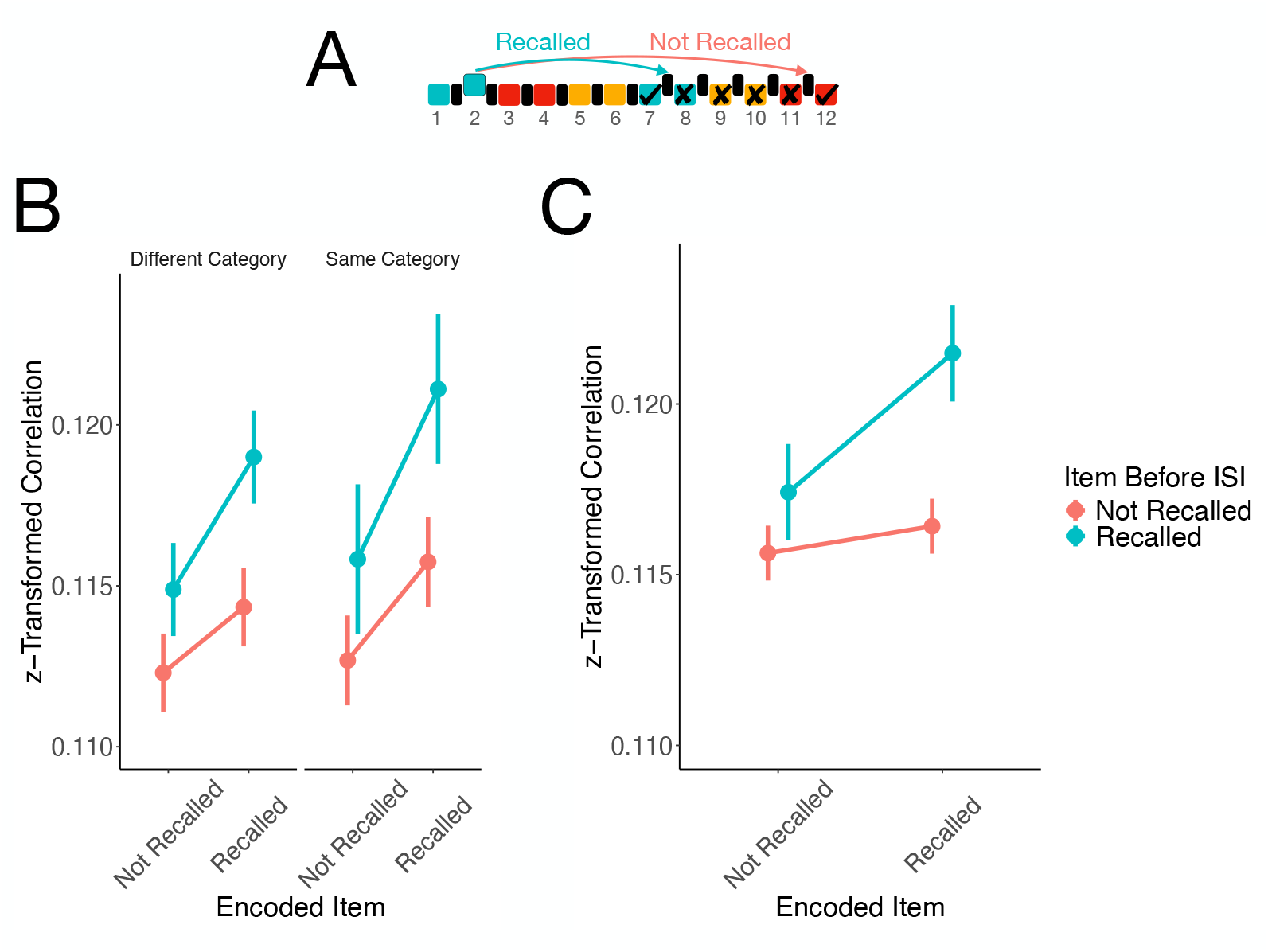
Similarity of neural signals at time points during encoding word presentations and time points during inter-item intervals as a function of whether the word was itself subsequently recalled. Estimates represent marginal means derived from the mixed effect model described in the main text. Error bars reflect 95% confidence intervals on the difference between remembered and forgotten, using an algorithm implemented by the R package `emmeans` and described in [41] that is a generalization of Loftus-Masson intervals [42, 43]. A. Schematic showing time points being compared when computing neural similarity. Specifically, we compare encoding time points from first half of list and inter-stimulus intervals from second half. Checks and crosses indicate whether the item was subsequently recalled and colors indicate category, indicating the relevant control variables. B. results from model fit to all *N* =215 subjects in Experiment 1. C. results from model fit to all *N* =257 subjects in Experiment 2.

**Fig. 4.**
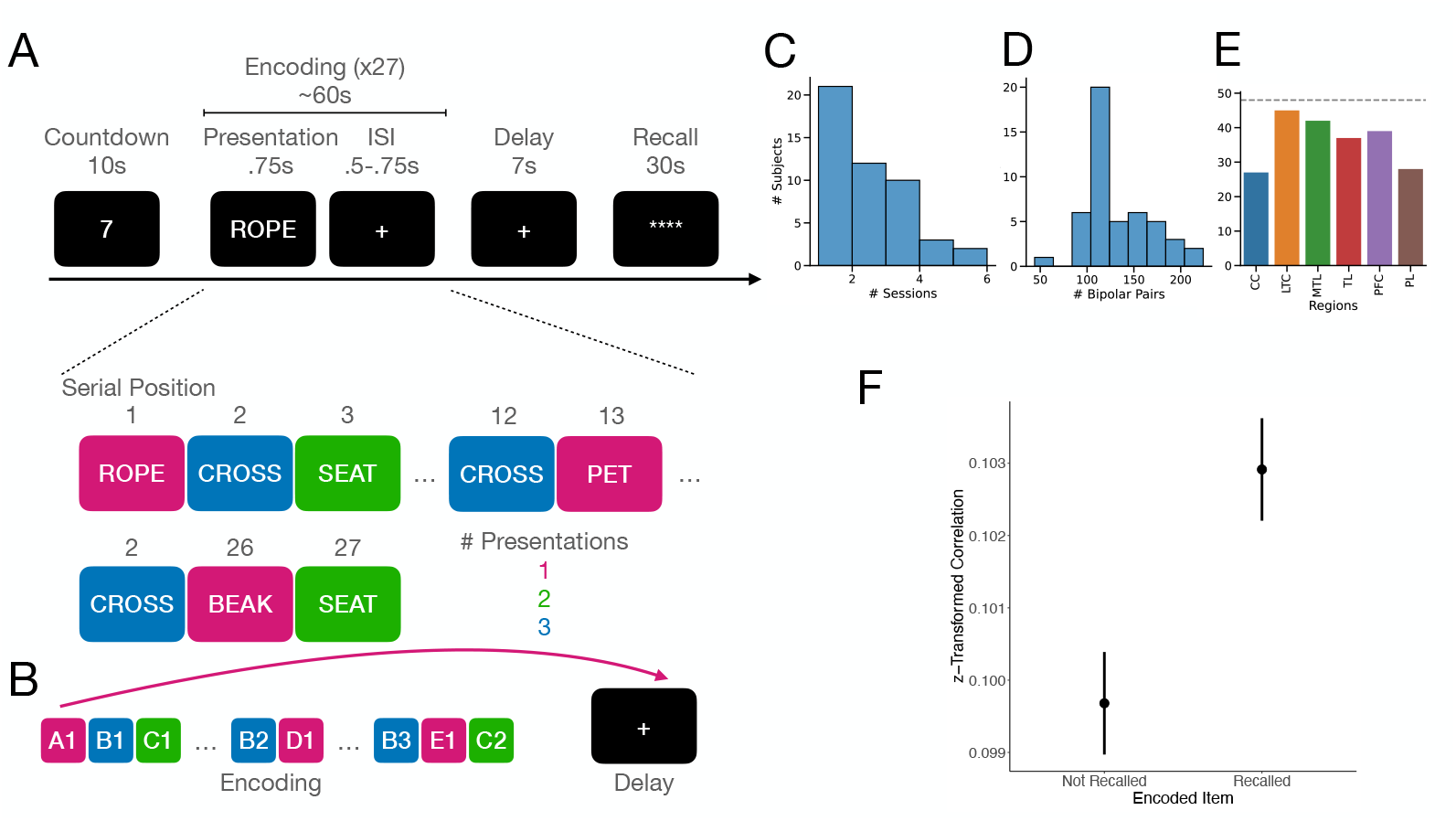
A. Task design for Experiment 3. Task is identical to Experiments 1 and 2 except that 1) lists include repeated items (some of which are presented twice and some presented three times) and 2) the math distractor task is replaced with an unfilled seven second delay interval. In addition, the items are presented on screen for shorter amount of time due to a bug in the experiment code. B. Schematic of comparisons used in analysis in panel F. Specifically, we compare neural activity during item presentations to neural activity during the delay interval. C. Distribution of sessions across subjects. D. Number of bipolar pairs per subject E. Number of subjects with bipolar pairs in each memory-related region. Grey dotted line indicates total number of subjects. F. Similarity of time points during encoding word presentations and time points during a delay period as a function of whether the word was itself subsequently recalled. Estimates represent marginal means derived from the mixed effect model described in the main text fit to all *N* =48 subjects. Error bars reflect 95% confidence intervals on the difference between remembered and forgotten, using an algorithm implemented by the R package `emmeans` and described in [41] that is a generalization of Loftus-Masson intervals [42, 43].

We fit the same model to Experiment 2, dropping the terms involving `same category`. We replicated the main effect of subsequent recall on study-phase spectral pattern similarity, i.e. items with more similar activity during subsequent ISIs during encoding than items that were not subsequently recalled (Fig. 3C, *F* (1, 14.22) = 5.98, *p* = .03). Investigation of the point estimates suggests that there is a positive difference between recalled and not-recalled. Additionally, the interaction between recall status of the encoding item and the item before the ISI was only marginally significant, (*F* (1, 11.81) = 4.02, *p* = .07), suggesting that we cannot reliably determine whether any specific pairwise comparison is different from the overall main effect. Again, we could not establish the direction of effects of first recall (*F* (1, 54.85) = .93, *p* = .34) or being a recall neighbor (*F* (1, 7.49) = .94, *p* = .36) on pattern similarity. Replicating results from Experiment 1, we find a main effect of the recall status of the item before the ISI (*F* (1, 39.86) = 7.48, *p* = .009).

To gain a more mechanistic understanding of these results, we asked whether our results were driven by activity in specific frequency bands. In the results just described, we computed correlations across eight frequency bands. We therefore repeated the statistical analysis while dropping a single frequency band from the similarity computation. This approach preserves the possibility that correlations were driven by across-frequency relationships between spectral signals. We found that the results were not particularly sensitive to any one frequency band being removed (Fig. A1) suggesting that the measured reinstatement is occurring generally at all spectral frequencies. We might be concerned that the analysis approach is more sensitive to variation in lower frequency bands because they do not have the opportunity to vary as much over the encoding interval. However, we don’t see any consistent trends across the frequency spectrum in the magnitude of results, suggesting that our results were not driven by any particular range of frequencies.

Perception of items often doesn’t require the full 1600 ms stimulus presentation interval. If subjects rapidly switch between online and offline states, it is possible that subjects switch into an offline state even while other items are on the screen. We therefore investigated whether reinstatement may be happening during the presentation of other items on the list. We replicated our analyses above using the last 500 ms of the 1600 ms stimulus presentation interval as the reinstatement period. We find that the main effect of recall status replicates across both experiments (Fig. B2). While not all pairwise comparisons are significant, they are all in the same positive direction as the results during the ISIs. It is to be expected that effects will be slightly weaker here as the signal should be dominated by the processing and encoding of the item currently on the screen. As a whole, these results suggest that switches between encoding and retrieval states may be happening continuously throughout the task phases.

### 2.3 Post-encoding reinstatement

Having identified study-phase reinstatement in two experiments, we sought to confirm that similar methods could replicate previous findings of post-encoding reinstatement [2, 3, 44] in the context of free recall of word lists. Therefore, in a third experiment, we investigated pattern similarity between initial encoding presentations and a subsequent unfilled 7s delay interval prior to recall. To test differences statistically, we fit the following statistical model:

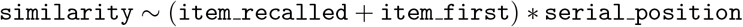

Including interactions with serial position allows for testing the possibility that earlier serial positions showed greater effects of reinstatement or that distributed reinstatement resulted in greater effects on recall than massed, as suggested by the literature on overt rehearsal [15, 17]. As in the analyses of the inter-stimulus intervals, we find a main effect of recall status such that remembered items are more similar to the delay interval than forgotten items (*F* (1, 135.87) = 43.16, *p* < .001. We additionally find a main effect of serial position (*F* (1, 6 9.03) = 86.35, *p* <. 001) but do not find any interactions between the recall effect and serial position (*F* (1, 30.06) = .48, *p* = .49). Examination of the point estimates suggests that similarity increases with later serial positions, likely due to auto-correlated noise in the signal. In this analysis, we are able to detect an effect of first recalls compared to later recalls (*F* (1, 23.44) = 5.28, *p* = .03) but the interaction with serial position was not statistically reliable (*F* (1, 17.91) = 1.36, *p* = .26). Examination of pairwise comparisons suggest that first recalls are actually less similar to delay intervals than other recalls, even with the adjustment for serial position.

To more closely relate our findings to those of Staresina and colleagues [2], we measured memory-related reinstatement during the math distractor tasks in Experiments 1 and 2. We replicate the main effect of recall status on reinstatement during the post-encoding interval in Experiment 3 (Fig. C3). Studies of overt rehearsal suggest that early list items show greater benefit of rehearsal [15] and that spaced rehearsals are more effective than massed [17]. Interestingly, we find an interaction with serial position but in the opposite direction predicted by overt rehearsal, such that the difference between remembered and forgotten items is greater for items towards the end of the list, closer in time to the delay interval.

## 3 Discussion

Three intracranial EEG experiments reveal that subsequently recalled items exhibit greater reinstatement during unfilled intervals than forgotten items. In the first two experiments, these intervals occurred between study opportunities, a time period that figures prominently in psychological theorizing on rehearsal but that has been overlooked in studies of neural reinstatement. During these interstitial periods, neural reinstatement appeared to be item- and list-specific. Our findings demonstrate a striking relation between spontaneous, sub-second, study-phase reinstatement and subsequent recall.

Psychologists frequently use study-phase retrieval as an explanatory construct [10– 14] for why rehearsal, repetitions, repeated testing or even presentations of similar items can enhance memory. While theoretically compelling, direct evidence for study-phase retrieval remains elusive because of the lack of corresponding overt behavioral correlates. Here, we provide evidence of a potential neural foundation for such retrieval processes, bolstering such claims and also allowing future work to investigate specific predictions in more detail. Our results also broaden the scope of previous work on neural reactivation and its effects on subsequent memory to include time periods during encoding events, rather than only offline rest periods.

We sought to identify evidence of spontaneous reactivation but activity driven by the item prior to the inter-stimulus interval carrying over into the delay period introduces potential confounds in analyses of study-phase reinstatement. Sepcifically, this activity could be similar to prior items for two reasons unrelated to spontaneous reactivation. One reason is that semantically similar items will have more similar neural representations than different ones [45]. Semantic similarity effects [46] imply that the probability of recalling an item increases when semantically similar items are also recalled. Thus, strong semantic representations of the item prior to the ISI could predict greater recall probabilities. A second reason is that similar cognitive processes are involved in processing the item. Subsequent memory effects [47–49] show that neural activity measured during encoding predicts subsequent recall. Thus, the similarity of neural responses to an item and other recalled items on a list should predict the item’s probability of recall. To address these confounds, we include terms in the model for whether the item before the ISI was itself recalled and, in the case of Experiment 1, whether it is from the same category as the earlier item. We did not detect a reliable interaction between the recall status of the encoded item and the recall status of the item before the ISI in either experiment. In addition, all pairwise comparisons point in the same direction and, in experiment 1, we find significant or marginally significant pairwise differences in all conditions, even in the case where the item before the ISI was not recalled and from a different category. Thus, we conclude that our findings likely reflect spontaneous reactivation.

Does our demonstration of spontaneous reinstatement reflect controlled rehearsal processes or a more unconscious and automatic process, such as that envisioned by systems consolidation theories [50, 51]? Our analyses of the math distractor intervals in Experiments 1 and 2 (Fig. C3), replicating findings from Staresina and colleagues [2] suggest that the measured reinstatement is likely, at least in part, automatic. Overt rehearsal studies show that when subjects rehearse items together they also recall them together and subjects tend to recall items with more rehearsals earlier in output [15]. In addition, the effectiveness of rehearsals on memory depends on spacing [17]. To the extent that our participants/subjects engaged in active rehearsal, and that our measure of neural reinstatement tracks such rehearsals, we can make three key predictions. First, finding greater study-phase reinstatement of an item from an earlier list position in the inter-stimulus interval immediately following a later item should increase the likelihood of those two items being successively recalled (as compared with being recalled in non-sequential output positions). Second, items with greater reinstatement should be more likely to be recalled first. Third, reinstatement further away in time from initial presentation should have a greater impact. However, we did not find evidence for any of these effects. This could indicate a difference between overt and covert rehearsal, with our measure of neural similarity being more sensitive to the latter. Alternatively, detecting these predicted organizational effects may require greater precision and thus more statistical power than detecting overall recall effects that integrate data across all of the interstimulus intervals.

One might also ask whether the identified reinstatement reflects specific content or the context associated with a retrieved episode [13]. Studies of reinstatement during delay periods [1–3], including the present one, have focused on neural similarity between a specific episode and a subsequent delay. Manning and colleagues [36] identified a neural contiguity effect during retrieval such that neural activity prior to retrieval is similar to items nearby the to-be-retrieved item. Howard and colleagues [52] observed similar effects in response to repeated items in a continuous recognition task. Inspired by models of memory [53, 54], these papers have claimed that this is evidence for reinstatement of the surrounding context. This literature has largely focused on retrieval and we do not know of any papers investigating the relationship between reinstatement of context during a delay interval and subsequent memory (although perhaps the work by Wu and colleagues [32, 33] could be interpreted in this light). Without analyzing the relationship between recall of a specific item and neural similarity of delay intervals to its neighbors, it is impossible to differentiate these two accounts of neural reinstatement. We therefore leave this important question to future work.

Our study focused on brain-wide reinstatement of multivariate neural activity and was thus not tailored to addressing questions about the specificity of these effects to specific neural regions. Moreover, intracranial EEG recordings provide high spatial and temporal precision but there is frequently little overlap in electrode locations across patients making it difficult to investigate anatomical substrates of multivariate signals and their consistency across people. We therefore leave investigation of anatomical specificity to future work, with more regionally-focused data sets or using recording methodologies that allow for broad coverage and alignment across subjects (e.g. fMRI).

In this work, we focused on a single memory task with relatively short study-test delays. This allowed us to easily test generalization across variants of this task (with surprisingly close quantitative estimates in figures 3B and C) but one may ask whether the effects here generalize to the longer time scales investigated in studies involving sleep [3, 4]. Although we demonstrate covert reactivation within the short interstimulus intervals and we measure recall following a 30 second filled distractor period, the effects of study-phase reactivation may be different after hours or days. Here we propose that study-phase reinstatement is a consolidation-like mechanism but most studies of consolidation [55, 56] suggest it is a gradual process with a time scale on the order of days to weeks rather than seconds as in the current study. We recognize that other mechanisms may operate at longer time scales but propose two ways in which the mechanisms might be related. First, due to the recursive nature of memory, as proposed by retrieved context theories [54], study-phase retrieval events that enhance memory on the scale of minutes (as in the present study) will likely lead to future reinstatement events that will further enhance memory. Such reinstatements may occur during quiescent periods and even during sleep, leading to consolidation effects at very long time scales. Another possibility, pointed out by an astute reviewer, is related to recent findings that the use of semantic schemas can promote much more rapid consolidation [57] and the fact that the most effective form of rehearsal is elaborative [58], involving deep semantic processing. This suggests that perhaps semantic processing is a key mechanism that allows for rapid consolidation. Consolidation may also may operate through multiple mechanisms on different time scales [20]. Future research examining the relation between sub-second covert reactivation and retrieval after longer delays will be required to obtain a more complete understanding.

The present findings show that covertly and spontaneously reactivated neural traces while encoding other experiences predict future memory. Overall, this work contributes to the greater cognitive neuroscience literature on memory consolidation in three ways. First, the ability to observe the memorial consequences of study-phase reinstatement suggests that memory consolidation is a much more general phenomenon than many previously thought. It likely occurs throughout awake experience with rapid switches between online, externally-focused, encoding states and offline, internally-focused, consolidation states. We bring human intracranial recordings to bear on the question of whether there is a special role for longer offline intervals such as sleep [6, 7, 22] in consolidation and find support for recent theories suggesting a more opportunistic view [20, 21, 24]. Second, we provide direct neuroscientific evidence of spontaneous study-phase retrieval, a cognitive mechanism theorized based on behavioral phenomena but lacking biological data. Finally, we compare stylized facts about overt rehearsal, a proposed psychological mechanism for rapid consolidation, to neural reinstatement. We fail to find consistent evidence that neural reinstatement relates memory organization in the same way as rehearsal, even with multiple highly-powered datasets (by the standards of the extant neuroimaging literature). We believe these contributions provide important constraints on theories of memory consolidation, a fundamental area of inquiry in the cognitive neurosciences.

Many further open questions remain about the psychological and neural mechanisms. To what extent are the reinstatement processes under cognitive control? Are the physiological mechanisms are the same as consolidation during sleep and over longer time scales? With the ability to directly measure these processes with neural data, we can now begin to answer these questions and many others. Identifying the connection between neural consolidation processes and psychological constructs developed to explain behavioral data will allow for a tighter link in theory development across cognitive and neural levels of analysis, a central goal of the cognitive neuroscience of memory.

## 4 Methods

### 4.1 Task and participants

In Experiment 1, we recorded intracranial EEG from 215 neurosurgical patients while they performed a categorized free-recall task [59]. Lists consisted of 12 items, presented in same-category pairs, from three distinct categories with categories drawn from total set of 25. Each item was on the screen for 1600 ms followed by an inter-stimulus interval with the length randomly jittered across trials. The interval on each trial was drawn uniformly on an range with a minimum of 750ms and a maximum of between 1000 and 1180 ms depending on which version of the experiment was used. Presentation of words during encoding was followed by a 20s post-encoding delay, during which subjects performed an arithmetic task in order to disrupt memory for end-of-list items (Fig. 2). Finally, subjects attempted to freely recall as many words as possible during a 30 second interval.

In Experiment 2, we collected EEG from 257 neurosurgical patients while they performed a standard free-recall task. Lists consisted of 12 items selected so that the average pairwise semantic similarity based on latent semantic analysis [60, 61] was .2. Timing of the item presentations, inter-stimulus intervals, distractor task and recall period were the same as in Experiment 1.

In Experiment 3, we collected EEG from 48 neurosurgical patients while they performed a free-recall task with repeated items. Lists consisted of 12 unique items and used the same wordpool as Experiment 2. However, 3 items were repeated twice and six items were repeated three times resulting in twenty-seven total encoding presentations per list. For the first 16 subjects items were on the screen for 1600 ms followed by an inter-stimulus interval of 750-1000ms, with the length of the interval randomly jittered across trials. Due to a coding error, the final 33 subjects only saw items on the screen for 750ms. Presentation of words during encoding was followed by a 7s unfilled post-encoding delay. As the previous experiments, subjects then attempted to freely recall as many words as possible during a 30 second interval.

All tasks were programmed using custom code with PyEPL [62] and UnityEPL. Our research protocol was approved by the Institutional Review Board at each participating university prior to data collection for all experiments, including Dartmouth Health Human Research Protection Program at Dartmouth-Hitchcock Medical Center (Hanover, NH), the Emory Human Research Protection Program at Emory University Hospital (Atlanta, Georgia), Hospital of the University of Pennsylvania (Philadelphia, PA), Mayo Clinic (Rochester, MN), Thomas Jefferson University Hospital (Philadelphia, PA), Columbia University Medial Center (New York, NY), and University of Texas Southwestern Medical Center (Dallas, TX). Additionally, informed consent was obtained from the participants and their guardians.

### 4.2 Intracranial EEG

As described in the original presentation by Weidemann and colleagues [59], we recorded from subdural grids and strips (space between adjacent contacts: 10 mm) and from depth electrodes (space between adjacent contacts: 5–10 mm) on a variety of recording systems across clinical sites with sampling rates varying between 500 and 2000 Hz to accommodate the local recording environment. We rereferenced all recordings using a bipolar referencing scheme [63] and applied a 4th order Butterworth filter with a 58-62 Hz stop-band to remove line noise. We then Fourier resampled the data to 1000 Hz and convolved the resulting signals with Morlet wavelets (wave number 5; 8 center frequencies, log-spaced between 3 and 180 Hz) to obtain a representation of spectral power in each bipolar pair. When computing power, we always include a buffer of 1 second on either side of the period of interest. This ensures that there are sufficient samples for estimating power in all frequency bands of interest. For the analysis of encoding data, we averaged power over the entire 1600 ms stimulus presentation interval (we use 15-1575ms following stimulus onset to avoid effects of luminance changes). We computed this average prior to running the similarity analyses. In Experiment 3, due to the coding error mentioned in the task description, we only used the first 750ms of the item presentation interval. For analysis of the inter-stimulus intervals, we use the same approach but for the 700ms interval immediately preceding the presentation of the subsequent stimulus. Similar to the approach for encoding periods, we compute this average prior to running the similarity analyses. For the analysis of the delay interval data in Experiment 3, we first preprocess all 7 seconds using the same approach as above. We then divide it into 7 one second bins and average over the 1 second, using a resampling technique (implemented in scipy [64]). For analyses presented here, we then average over the 1 second bins. We compute the similarity analyses within each 1 second bin and then average the 7 bins to obtain a single similarity value for each encoded item. For analysis of the end of the encoding interval in Appendix B, we followed the same approach as for the inter-stimulus interval but using the last 500ms of the stimulus presentation. For analysis of the math distractor task data from Experiments 1 and 2 in Appendix C, we followed the same approach by dividing up the interval in 20 one second bins and then averaging.

In order to compare activity across periods and subjects, we normalize all data relative to spectral signals collected during the 10 second countdown period prior to each list. We first preprocess the 10 seconds using the same approach as above. We then divide it into 10 one second bins and average over the 1 second, using the resampling technique mentioned above. We then compute the mean and standard deviation across lists for each session. All spectral power estimates in subsequent periods are then normalized by subtracting the countdown mean and dividing by the countdown standard deviation.

### 4.3 Analysis

Our main analysis approach relies on analyzing the similarity of spectral power patterns [34, 35] in the eight frequency bands across all bipolar pairs, regardless of location in the brain. As is common in the literature, similarity was determined as the Fisher’s *z*-transformed Pearson correlation [38] between power at two time points. These analyses relied on `xarray` [65], `pandas` [66], `scipy` [64] and `numpy` [67] in `python 3.11`. For visualization we also used `seaborn` [68].

### 4.4 Modeling

For all models, we use mixed effect models to account for systematic variation unrelated to our effects of interest, as well as variability in these effects across the population and task structure. We fit the models using `lme4` [69] and `lmerTest` [70], with *p* values for all inferential *t* and *F* tests determined using the Satterthwaite approximation [71–73].

In our first model, we test whether pattern similarity between two word presentations at encoding is driven by both semantic and temporal distance. We allow these effects to vary by subject, session, list and serial position and their interactions. The full model that we fit, as specified in `lme4`, is:

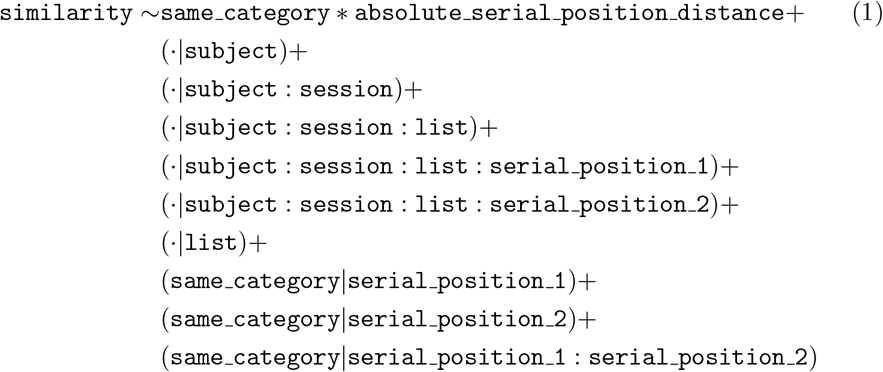

where `same category` indicates that the two items are from the same category and `absolute serial position distance` indicates the distance in terms of serial positions between the two items as a linear effect. The dot (·) indicates that all terms vary as a function of that random effect grouping variable. `list` indicates the list position within the session (from 1 to 25) and `serial position 1` indicates the earlier serial position in the pair while `serial position 2` indicates the later one. For this analysis we used all possible combinations of earlier and later serial positions. Because the large number of random effects could lead to overfitting, we reduce the model iteratively to account for the structure present in the data using the procedure from [74]. We use the same procedure for all subsequently described models as well.

In the second set of models, we test whether pattern similarity between a word presentation and a later inter-stimulus interval at encoding is driven by whether the initial word was recalled, whether it was recalled first and whether it was recalled next to the item before the ISI. For this analysis, we only used word presentations in the first half of the list and inter-stimulus intervals in the latter half. This ensured that each encoded item tested could be reinstated during each inter-stimulus interval. We also adjust for potential effects and interactions with semantic similarity and whether the item before the ISI was itself recalled. We again allow these effects to vary by subject, session, list and serial position. In addition, the model included random effects for across-subject effects of list ID, serial positions of the encoded item and the ISI and the interaction between the pair of serial positions. The full model that we fit, as specified in `lme4`, is:

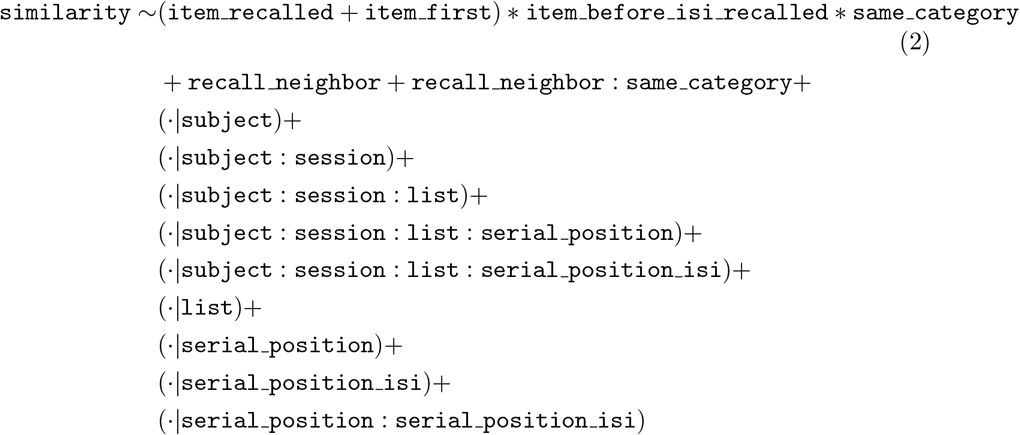

`item recalled` and `item first` reflect whether the specific item was recalled and whether it was recalled first. `item before isi recalled` indicates whether the item before the isi was recalled and `same category` indicates that the initial encoding item and the item before the ISI are from the same category. Finally, `recall neighbor` indicates that, if both items were recalled, whether they were recalled next to each other during the recall period. The dot (·) indicates that all terms vary as a function of that random effect grouping variable. `serial position` indicates the serial position of the initial presentation while `serial position isi` indicates serial position preceding the ISI.

In the final set of models, we examine the degree to which items that are subsequently recalled are reinstated during a delay interval between the encoding phase and the test phase. To test this, allowing for potential interactions with serial position, we use the following specification:

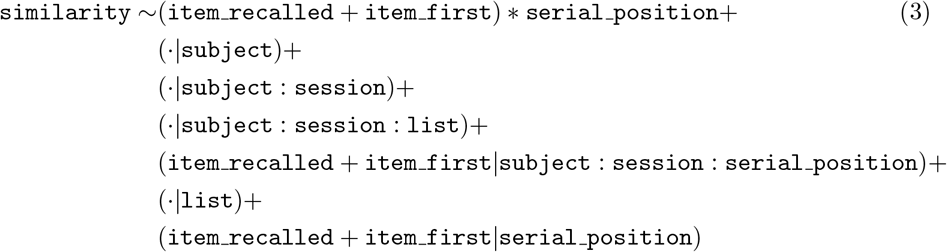

This model captures serial position effects with a continuous fixed effect as well as random effects for each serial position that allow for deviations from linear effects.

For this analysis, we use all encoded items, because they all have the potential to be reinstated during the delay.

Post-hoc analysis of models relied on `tidyverse` [75] and `emmeans` [41] in `R 4.3.2` [76].

## Acknowledgements

We are very grateful to all patients who participated in this study. We thank R. Colyer, A. Rao, and R. DeHaan for help with data collection and quality control and A. Schapiro, A. Broitman and N. Greene for comments on the manuscript. M.J.K. was supported by NIH grant MH55687

## Competing interests

The authors declare no competing interests.

## Availability of data and materials

De-identified data from the three experiments in this study are available on OpenNeuro (Experiment 1: https://openneuro.org/datasets/ds004809/, Experiment 2: https://openneuro.org/datasets/ds004789/, Experiment 3: https://openneuro.org/datasets/ds005411/)

## Code availability

The software used to perform analyses and generate figures for this manuscript is available on GitHub at https://github.com/pennmem/study_phase_reinstatement.

## Authors’ contributions

D.J.H. and M.J.K. developed hypotheses and designed the study. M.J.K. acquired funding. B.C.L., R.E.G., C.W., M.R.S., J.P.A., B.C.J. recruited study participants, implanted electrodes and collected the data. D.J.H. analyzed the data. D.J.H. and M.J.K. wrote the paper.

## Appendix A Frequency Specificity

In order to obtain a more mechanistic understanding of the frequencies contributing to our observed rapid reinstatement results, we replicated our analysis dropping individual frequency bands when computing the correlation between timepoints. Fig. A1 replicates Fig. 3, showing the estimated difference in similarity to subsequent ISIs when the encoding trial is recalled vs. not recalled when each of the eight frequency bands are dropped in each subgroup of trials. The results indicate that the effect is largely unchanged when individual frequencies are dropped, suggesting that all frequencies contribute to the observed reinstatement.

**Fig. A1.**
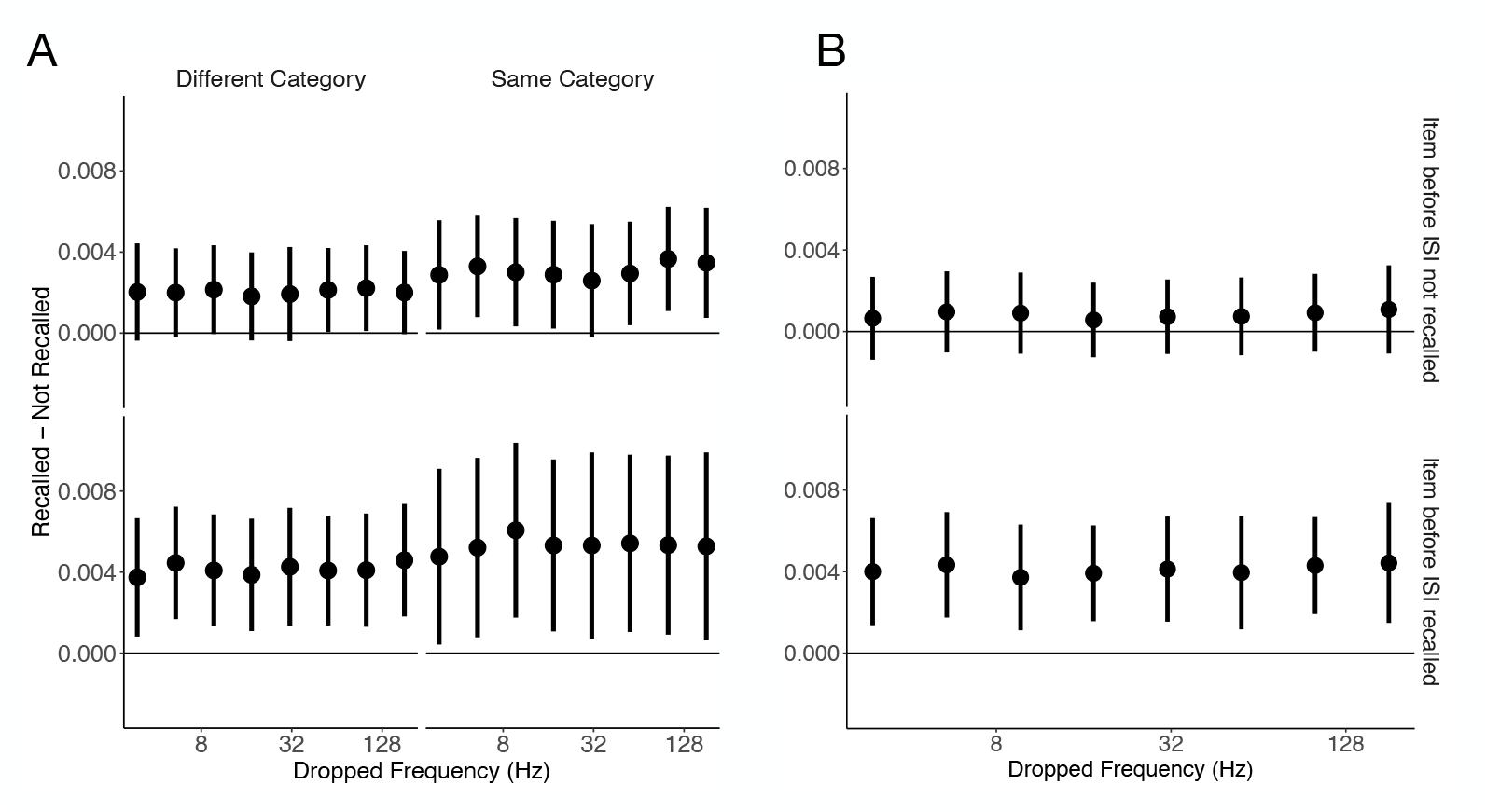
Replication of Fig. 3 while dropping individual frequency bands from the similarity computation. X axis indicates which frequency is dropped and is shown on a log scale. A. results from models fit to all *N* =215 subjects in Experiment 1. B. results from models fit to all *N* =257 subjects in Experiment 2.

## Appendix B Generalization to end of encoding presentation

In Experiment 1, we replicate the results from our ISI analysis, finding a main effect of `item recalled` such that items that were subsequently recalled were more similar to activity during the last 500ms of subsequent encoding items items that were not subsequently recalled (Fig. B2, *F* (1, 8.3) = 2.77, *p* = .02). However, we could not reliably identify whether the items recalled first had consistently different levels of reactivation than other recalled items (*F* (1, 127.8) = .004, *p* = .95). We also did not see consistent differences between items that were sequentially recalled and pairs of items that were both recalled but not sequentially (*F* (1, 9.3) = 3.93, *p* = .08). As with the ISIs, we find that the recall status of the currently encoded item predicts similarity (*F* (1, 37.2) = 31.43, *p* < .001). We also find that the two items being from the same category is a significant predictor (*F* (1, 272.2) = 37.2, *p* = .03). Unlike above, we find that the recalled effect differed depending on whether the current item was from the same category or not (*F* (1, 16727.2) = 8.18, *p* = .004). Inspection of Fig. B2 suggests that this is due to the effect being larger when the two items are from the same category. Despite this, all estimates are in the same positive direction, suggesting that this is picking up on a similar effect to the ISI analysis.

We again fit the same model to Experiment 2, dropping the terms involving `same category`. We replicated the main effect of subsequent recall on study-phase spectral pattern similarity, i.e. items with more similar activity during subsequent ISIs during encoding than items that were not subsequently recalled (Fig. 3, *F* (1, 9.49) = 9.94, *p* = .01). Similar to the ISIs, we do not find reliable evidence for an interaction with the recall status of the current item (*F* (1, 19.75) = 3.75, *p* = .08). Again, we could not establish the direction of effects of first recall (*F* (1, 5.54) = .67, *p* = .45). Interestingly, here we find evidence for an effect of being neighboring recalls (*F* (1, 30.56) = 6.64, *p* = .02) on pattern similarity.

**Fig. B2.**
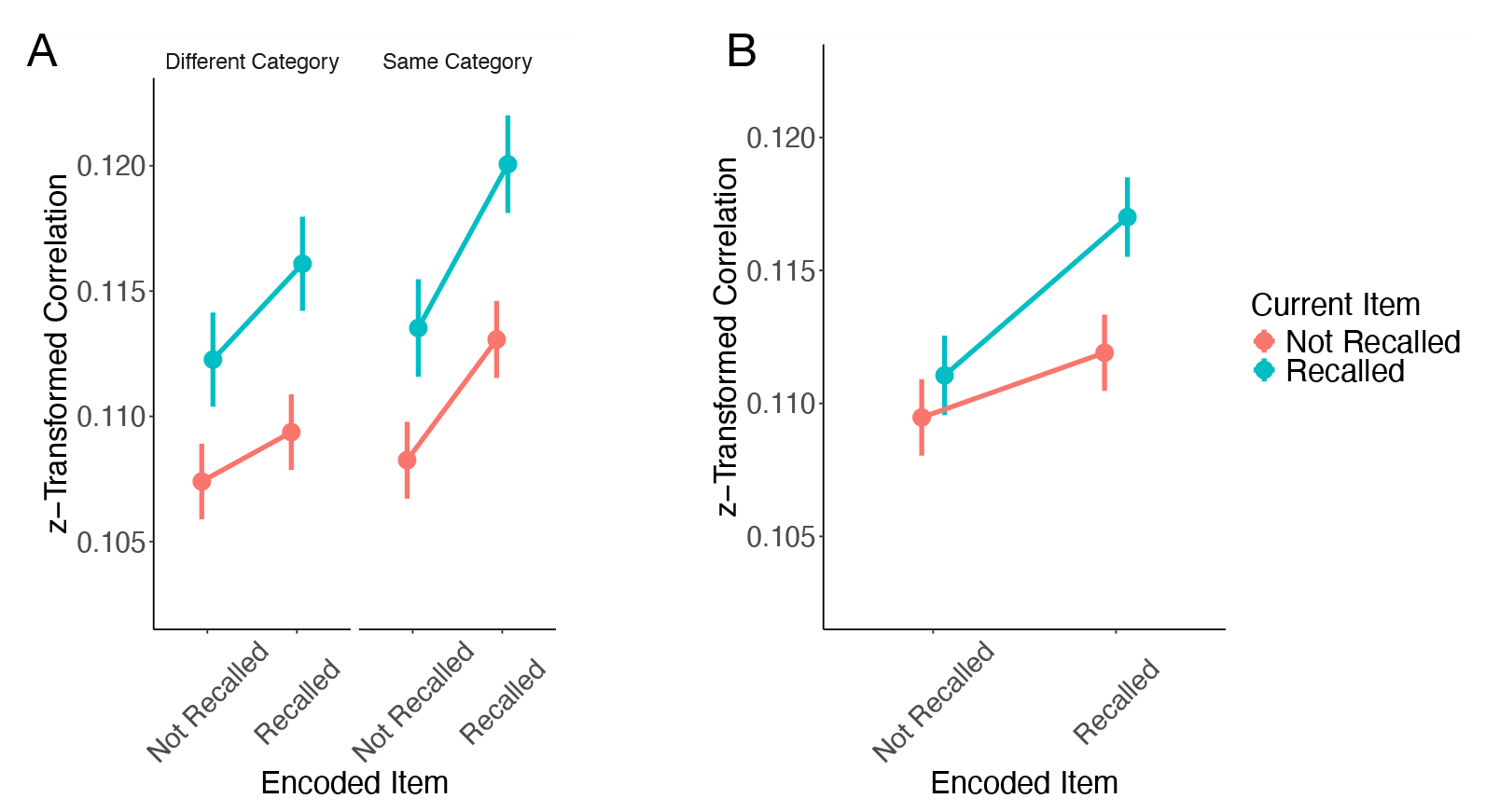
Similarity of time points during encoding word presentations and time points during the last 500ms of subsequent encoding word presentations as a function of whether the word was itself subsequently recalled. Estimates are marginal means derived from the mixed effect model described in the main text. Error bars reflect 95% confidence intervals on the difference between remembered and forgotten, using an algorithm implemented by the R package `emmeans` and described in [41] that is a generalization of Loftus-Masson intervals [42, 43]. A. results from model fit to all *N* =215 subjects in Experiment 1. B. results from model fit to all *N* =257 subjects in Experiment 2.

## Appendix C Generalization to distractor period

In order to more directly replicate findings from Staresina and colleagues [2] and investigate whether at least some of the measured reinstatement is likely to be unconscious, we analyzed data during the math distractor task in experiments 1 and 2 using the same model as used above in Experiment 3. We find a significant main effect of recall status on reinstatement (Experiment 1: *F* (1, 210.4) = 73.02, *p* < .001; Experiment 2: *F* (1, 236.59) = 87.79, *p ≤* .001). In both experiments, we also find a significant interaction with serial position (Experiment 1: *F* (1, 204.9) = 9.96, *p* = .001; Experiment 2: *F* (1, 42.2) = 4.73, *p* = .04). Inspection of the point estimates suggests that this is primarily due to the difference in reinstatement between remembered and forgotten items being larger in later serial positions. Finally, as expected, we find a main effect of serial position (Experiment 1: *F* (1, 119.2) = 56.01, *p* < .001; Experiment 2: *F* (1, 46.55) = 48.2, *p* < .001).

**Fig. C3.**
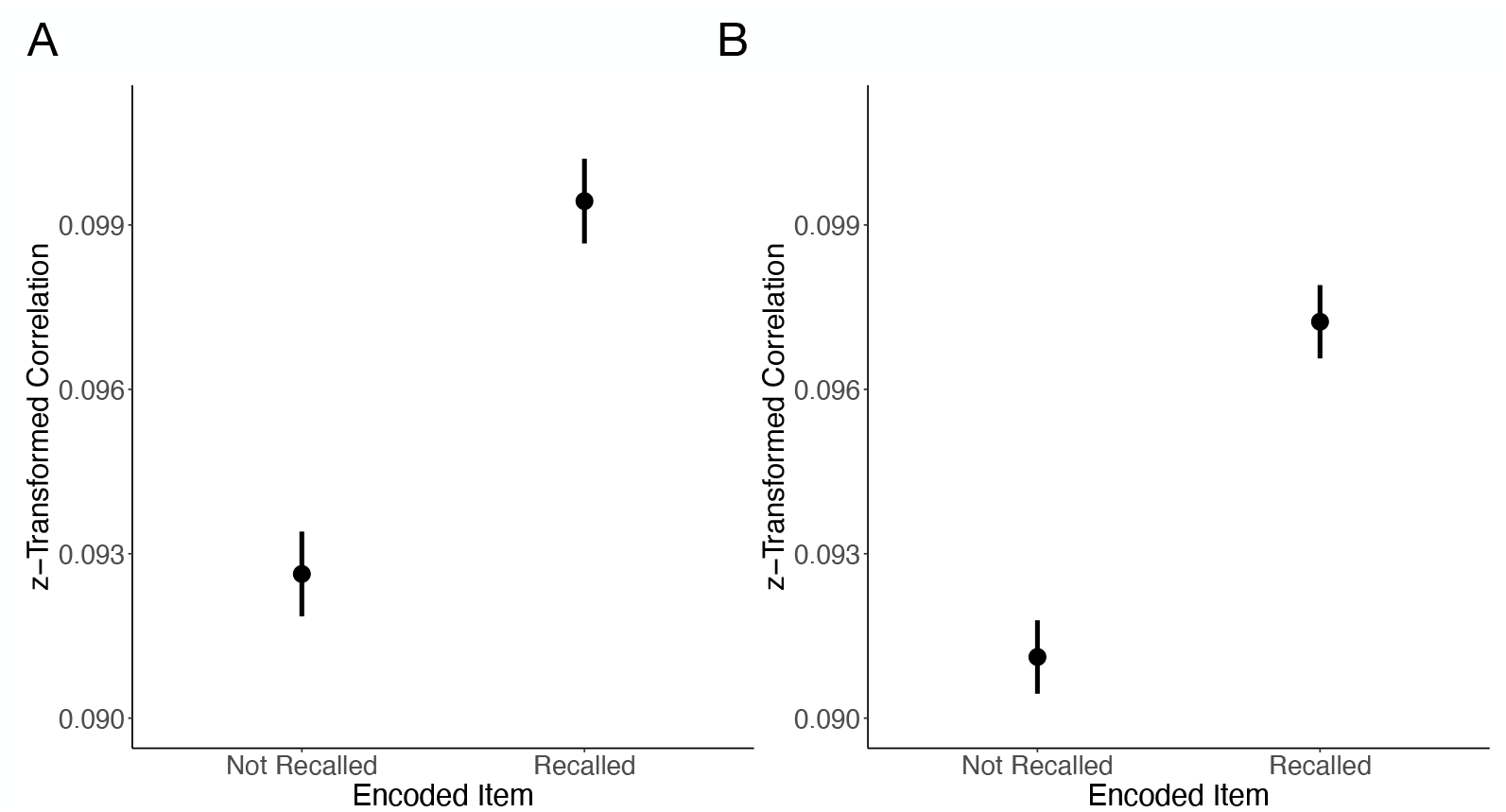
Similarity of time points during encoding word presentations and timepoints during the math distractor task as a function of whether the word was itself subsequently recalled. Estimates are marginal means derived from the mixed effect model described in the main text. Error bars reflect 95% confidence intervals on the difference between remembered and forgotten, using an algorithm implemented by the R package `emmeans` and described in [41] that is a generalization of Loftus-Masson intervals [42, 43]. A. results from model fit to all *N* =215 subjects in Experiment 1. B. results from model fit to all *N* =257 subjects in Experiment 2.

## Notes

### Competing Interest Statement

The authors have declared no competing interest.

### Summary of Updates

Various updates in response to journal reviews.

